# Inverse Potts model improves accuracy of phylogenetic profiling

**DOI:** 10.1101/2021.10.18.464903

**Authors:** Tsukasa Fukunaga, Wataru Iwasaki

## Abstract

Phylogenetic profiling is a powerful computational method for revealing the functions of function-unknown genes. Although conventional similarity evaluation measures in phylogenetic profiling showed high prediction accuracy, they have two estimation biases: an evolutionary bias and a spurious correlation bias. Existing studies have focused on the evolutionary bias, but the spurious correlation bias has not been analyzed. To eliminate the spurious correlation bias, we applied an evaluation measure based on the inverse Potts model (IPM) to phylogenetic profiling. We also proposed an evaluation measure to remove both the evolutionary and spurious correlation biases using the IPM. In an empirical dataset analysis, we demonstrated that these IPM-based evaluation measures improved the prediction performance of phylogenetic profiling. In addition, we found that the integration of several evaluation measures, including the IPM-based evaluation measures, had superior performance to a single evaluation measure. The source code is freely available at https://github.com/fukunagatsu/Ipm.

## Introduction

Genome sequences of many species have been determined, and accordingly, many function-unknown genes have been discovered. Revealing the functions of these function-unknown genes is an important research topic, but it is too time-consuming to experimentally verify the functions of all the genes. Therefore, the computational predictions of these gene functions are essential, and various bioinformatics methods have long been developed in bioinformatics. Phylogenetic profiling is one such analysis method. In this method, when two ortholog groups (OGs) have similar occurrence patterns among species in an ortholog table, the two OGs are presumed to be functionally related ([1, 2, 3, 4, 5]). Phylogenetic profiling has been widely used to estimate the functions of function-unknown genes in various phylogenetic groups from prokaryotes to eukaryotes ([6, 7]).

In conventional phylogenetic profiling, similarities in occurrence patterns between two OGs are directly calculated from an ortholog table as evaluation measures, such as mutual information (MI). These evaluation measures implicitly assume that the species included in the ortholog table are independent of each other. This assumption is incorrect as the species have evolutionary relationships. In other words, conventional evaluation measures introduce an evolutionary bias in the estimation. Therefore, evaluation measures that consider evolutionary information and are more accurate than conventional evaluation measures have been proposed ([8, 9, 10, 11, 3]).

Another possible estimation bias is the spurious correlation bias between the two OGs. In statistics, the spurious correlation means that two unrelated (or weakly related) variables appear to be strongly related due to the influence of confounding factors. For example, suppose there are functional relationships between OGs A and B and OGs A and C, but no (or weak) functional relationship between OGs B and C. In this case, OGs B and C can show a high MI by bypassing OG A, which is a confounding factor. Ignoring the possibility of spurious correlations should negatively influence the accuracy of the function predictions. However, the spurious correlation bias has not been considered in previous phylogenetic profiling studies.

MI is a local evaluation measure calculated from only two OG profiles, and the locality causes spurious correlations whose confounding factors are the other OGs. Therefore, we can eliminate these spurious correlation biases by using a global evaluation measure calculated from all OG profiles. The inverse Potts model (IPM), also called direct coupling analysis or evolutionary coupling ([12]), is an analysis method for calculating the global evaluation measure. The IPM has been applied to various biological data analyses, such as protein-protein interaction prediction ([13, 14]), protein structure prediction ([15, 16]), neural data analysis ([17, 18]), and genome-wide association studies ([19, 20]), and has improved prediction performance. Recently, Croce *et al*. identified physically interacting protein domain pairs by applying the IPM to tabular data whose rows and columns are species and protein domains ([21]). They revealed that the IPM could detect interacting domain pairs with higher accuracy than simple correlation coefficients. Their study was similar to phylogenetic profiling, but their goal was to predict domain-domain interactions and not to estimate gene functional associations.

In this study, we applied the IPM to phylogenetic profiling to accurately predict gene functions. We used direct information (DI) calculated based on the IPM as the global evaluation measure. We also calculated the DI with evolutionary information to eliminate both the evolutionary and spurious correlation biases. We investigated the performance of several evaluation measures in phylogenetic profiling, and verified that the IPM-based evaluation measures improved the accuracy of predicting gene functions. In addition, we found that the integration of several evaluation measures, including the IPM-based evaluation measures, has superior performance to a single evaluation measure.

## Methods

### Input data

Two settings were assumed in our study: standard and evolutionary settings. Under the standard setting, the input data for our method is an ortholog table *D*, which consists of *N* species and *L* OGs. *D*_*i, j*_ represents whether species *i* has OG *j* and takes either 0 or 1. Under the evolutionary setting, the input data for our method is an ortholog gain/loss table *D*, which consists of *N* phylogenetic tree branches and *L* OGs. Given a phylogenetic tree and an ortholog table, any method for reconstructing gene-content evolutionary history can be used to infer gene gain/losses on each branch of the tree. *D*_*i, j*_ represents whether the gain/loss events of OG *j* occurred at edge *i*. The value takes 0, 1, or 2, indicating that there are no gene gain/loss events, gene gain events, or gene loss events, respectively.

For the experiments in this study, we used three empirical datasets: archaea (domain), micrococcales (order), and fungi (kingdom) ([22]). The ortholog tables were prepared by preprocessing ortholog tables in the STRING database ([23]). Under the standard setting, we ignored gene copy number information and removed OGs that were shared by less than 10% or more than 90% of the species to reduce the computational time (note that the computational time of our method is proportional to the square of the number of OGs) to prepare *D*. The removed OGs were expected not to have significant impacts on the results because of their low information content. The archaea, micrococcales, and fungi datasets consisted of 151 species and 2,875 OGs, 111 species and 1,905 OGs, and 123 species and 5,786 OGs, respectively. Under the evolutionary setting, we prepared *D* by reconstructing the gene-content evolutionary history for the three empirical datasets. We used Mirage ([22]) with the BDARD model ([24]) and the PM model (default parameters were used for the others). Phylogenetic trees were supplied by the Genome Taxonomy Database release 89 ([25]) for the archaea and micrococcales datasets and the SILVA database release 111 ([26, 27]) for the fungi dataset.

### The IPM

We introduce MI_*ab*_, which is the MI between OG *a* and OG *b*. The formula is as follows:

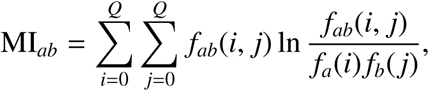

where *f*_*a*_(*i*) and *f*_*ab*_(*i, j*) are the relative frequencies of OG *a* taking *i* and OG *a* and OG *b* taking *i* and *j*, respectively, in the dataset *D. Q* is the maximum value that an OG can take (that is, *Q* = 1 under the standard setting and *Q* = 2 under the evolutionary setting). The more OGs A and B depend on each other, the larger the MI_*ab*_. If MI_*ab*_ becomes 0, OGs A and B are completely independent. We defined standard MI (SMI) and EMI as MI calculated under the standard and evolutionary settings, respectively.

MI_*ab*_ is a local evaluation measure calculated from only two OG profiles and is vulnerable to spurious correlations. Therefore, we calculated a global evaluation measure using all OG profiles based on the IPM. We first formulate the joint probabilities of all OGs as follows ([12]):

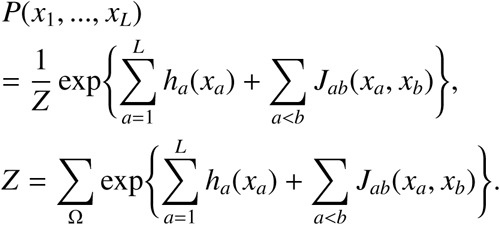

*P*(*x*_1_, …, *x*_*L*_) is the joint probability that OG *a* takes *x*_*a*_ for any *a. h*_*a*_(*x*_*a*_) is a weight parameter when OG *a* is *x*_*a*_, and *J*_*ab*_(*x*_*a*_, *x*_*b*_) is also a weight parameter when OG *a* is *x*_*a*_ and OG *b* is *x*_*b*_. Ω is the set of all possible combinations that all OGs can take, and *Z* is a normalizing constant, which is called the partition function. This probabilistic model is obtained by deriving a model that maximizes entropy under the following constraints: *f*_*a*_(*i*) = *p*_*a*_(*i*) for all *a* and *i* and *f*_*ab*_(*i, j*) = *p*_*ab*_(*i, j*) for all *a, b, i*, and *j. p*_*a*_(*i*) and *p*_*ab*_(*i, j*) are the marginal probabilities of *P*(*x*_1_, …, *x*_*L*_) and represent the probabilities of OG *a* taking *i* and OG *a* and OG *b* taking *i* and *j*, respectively. This model is generally called the Potts model in statistical physics (when *Q* = 1, this model is specifically called the Ising model). Note that this model is also a particular form of the Boltzmann machine or Markov random field.

To calculate the parameters *h*_*a*_(*i*) and *J*_*ab*_(*i, j*) analytically, we need to count all the combinations in Ω. However, its computational cost can become too large when *L* is large because the number of combinations becomes large. Therefore, these parameters are learned from the dataset in an unsupervised manner (Section 2.3). Then, using the estimated parameters, the dependence between OG *a* and OG *b* is measured as DI_*ab*_ as follows ([13]):

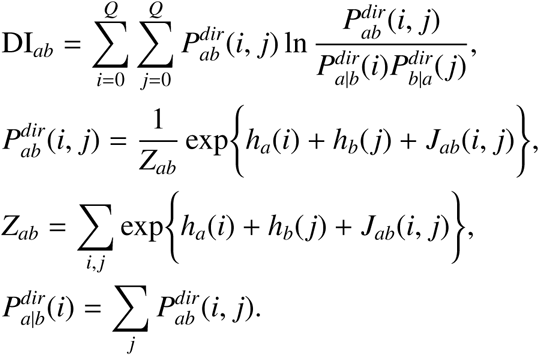

Note that this definition is slightly different from the original definition ([13]). In the original DI calculation, *f*_*a*_(*i*) was used instead of 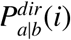, and *h*_*a*_(*i*) was recalculated from 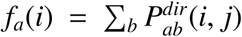. Similar to MI_*ab*_, the more OGs A and B depend on each other, the larger DI_*ab*_. We defined standard DI (SDI) and EDI as the DI calculated under the standard and evolutionary settings, respectively.

### Parameter estimation method

In the derivation of the Potts model, the number of substantial constraints is 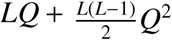 because ∑_*i*_ *f*_*a*_(*i*) = 1 and ∑_*i j*_ *f*_*ab*_(*i, j*) = 1 must be satisfied. On the other hand, the number of parameters in the model is 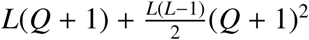, which is larger than the number of substantial constraints. This over-parameterization leads to the non-identification of the model. Therefore, it is necessary to introduce additional constraints on the parameters to reduce the degrees of freedom of the model. In this study, we used the following constraints, called lattice gas gauges ([12]):

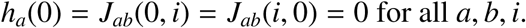

To date, various algorithms have been developed to estimate the parameters of the Potts model, for example, mean-field approximation ([28]), pseudo-likelihood maximization ([29]), adaptive cluster expansion([30]), and Markov chain Monte Carlo (MCMC) methods ([31]). There is an approximate trade-off between the computational speed and estimation accuracy in these methods, that is, more accurate methods require longer run times. In this study, we focused on the estimation accuracy, and used the persistent contrastive divergence (PCD) method ([32, 33]), which is a variant of the MCMC method. We maximized the likelihood with the L2-regularization term to avoid overfitting the data in the PCD method.

The algorithm for the PCD method is as follows. We first randomly sample *K* samples with replacement from the dataset *D*, and let the initial sampled dataset be *D*^0^. In this study, we set *K* to 200. In addition, we set all the initial parameters to 0. Next, we obtained the dataset *D*^1^ from *D*^0^ and the initial parameters based on the following Gibbs sampler:

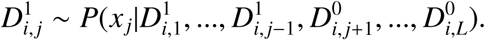

This sampling was performed *LK* times to obtain *D*^1^. Then, we calculated 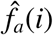 and 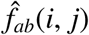, which are the relative frequencies of OG *a* taking *i* and OG *a* and OG *b* taking *i* and *j* in the dataset *D*^1^, respectively. Subsequently, the model parameters were updated using the following formula:

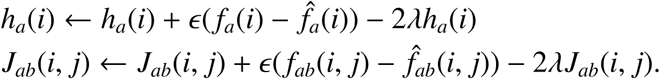

2*λh*_*a*_(*i*) and 2*λJ*_*ab*_(*i, j*) are the L2-regularization terms, and we used any 0, 0.01, 0.05, 0.1, 0.5, 1.0, or 5.0 as *λ*. Note that *λ* = 0 indicates simple likelihood maximization without the regularization terms. *ϵ* represents a learning rate, and we set either 0.01 or 0.001 as *ϵ*. After parameter estimation, we sampled dataset *D*^2^ from *D*^1^ using the estimated parameters. We finally adopted parameters after repeating the Gibbs sampling and the parameter update 3000 times.

### Evaluation method

We assessed the prediction performance of each evaluation measure using association scores between two OGs provided in the STRING database ([23]). The association scores in the STRING database were calculated by considering gene neighborhood conservation, gene fusion, co-expression, protein interaction experiments, other databases, text mining, and occurrence patterns. Because occurrence patterns should not be used in the assessment, we recalculated the association scores by ignoring the occurrence pattern similarities. If the recalculated association score of an OG pair was larger than the threshold *th*, we regarded the OG pair as positive data; otherwise, we regarded it as negative data. We used the threshold *th* from 0.5 to 0.9 in 0.1 increments. The sizes of each dataset are listed in Table S1. Note that the association scores of 0.7 and 0.9 are the lower limits of high and highest confidences, respectively, in the STRING database.

We first investigated the overall discrimination performance of each evaluation measure using the area under the receiver operating characteristic curve (AUC) scores. The AUC scores were calculated using the pROC R package ([34]). In addition, we assessed the prediction accuracy of the OG pairs that were highly ranked by each evaluation measure. Specifically, we defined the highly ranked OG pairs as the top *M* OG pairs in each evaluation measure, and calculated the positive predictive values (PPV) of these pairs (at *th* = 0.7). We used 100, 500, 1,000, 5,000, or 10,000 as *M*.

## Results

### Performances of single evaluation measures

We first assessed the overall discrimination performance of the four evaluation measures (SMI, EMI, SDI, and EDI) based on the AUC scores. We investigated 14 combinations of seven *λ* values and two *ϵ* values as IPM hyperparameters for calculating the SDI and EDI. In the following analyses, we used the hyperparameters showing the best AUC score for each dataset and each *th* value. The AUC scores are listed in Table S2-S7. Both hyperparameters had a large impact on the prediction performance. In addition, the optimal hyperparameters differed depending on the dataset and the *th* value. We also found that the optimal hyper-parameter *λ* was not 0.0 in many cases. This result means that L2-regularization was effective for achieving high discrimination performance.

We checked the distribution of each evaluation measure after normalizing the maximum value to 1.0, and calculated the skewness (Fig. S1-2). We found that the distribution was skewed to the right in all cases, that is, only a portion of OG pairs obtained high scores in each evaluation measure. In addition, we discovered that the consideration of both gene-content evolutionary history and usage of the IPM increases the skewness of the distribution. These results suggest that the evolutionary and spurious correlation biases were removed by the reconstruction of the gene content history and the IPM method.

Fig. 1A-C shows the results of the AUC analyses. We found that EMI outperformed SMI in all cases, which suggests that gene content history reconstruction is highly effective in phylogenetic profiling, which is consistent with previous studies ([9, 10, 3]). SDI was always better than SMI, except for one case where similar performances were obtained (*th* = 0.9 in the micrococcales dataset). These results also suggest that the consideration of spurious correlation is effective and that the IPM is valuable for removing biases. As expected, EDI showed the best performance in the archaea and micrococcales datasets, except for the same case where EMI and EDI showed comparable performances. On the other hand, SDI showed the best performance in the fungi dataset. This result indicates that, under the evolutionary setting of the fungi dataset, spurious correlations still may have functional information. In eukaryotic genome evolution, for example, due to the expansion of genome size, many genes can simultaneously increase their copy numbers. We interpreted such simultaneous gene gains as spurious correlations in EDI, even if they have true functional relationships. It should also be noted that the performance of EDI increased with increasing *th*, in contrast to the other cases. This may suggest that fungal gene pairs with high association scores are especially affected by these effects. The performance of EDI in the fungi dataset may also be caused by insufficient gene annotation. Although the recalculated STRING scores used gene neighborhood conservation and gene fusion, they are not effective in estimating eukaryotic protein functional relationships. We found that the proportion of positive data was much lower for the fungi dataset than for the other datasets (Table S1). This suggests that many functionally related OG pairs in nature were not given high association scores.

**Figure 1:**
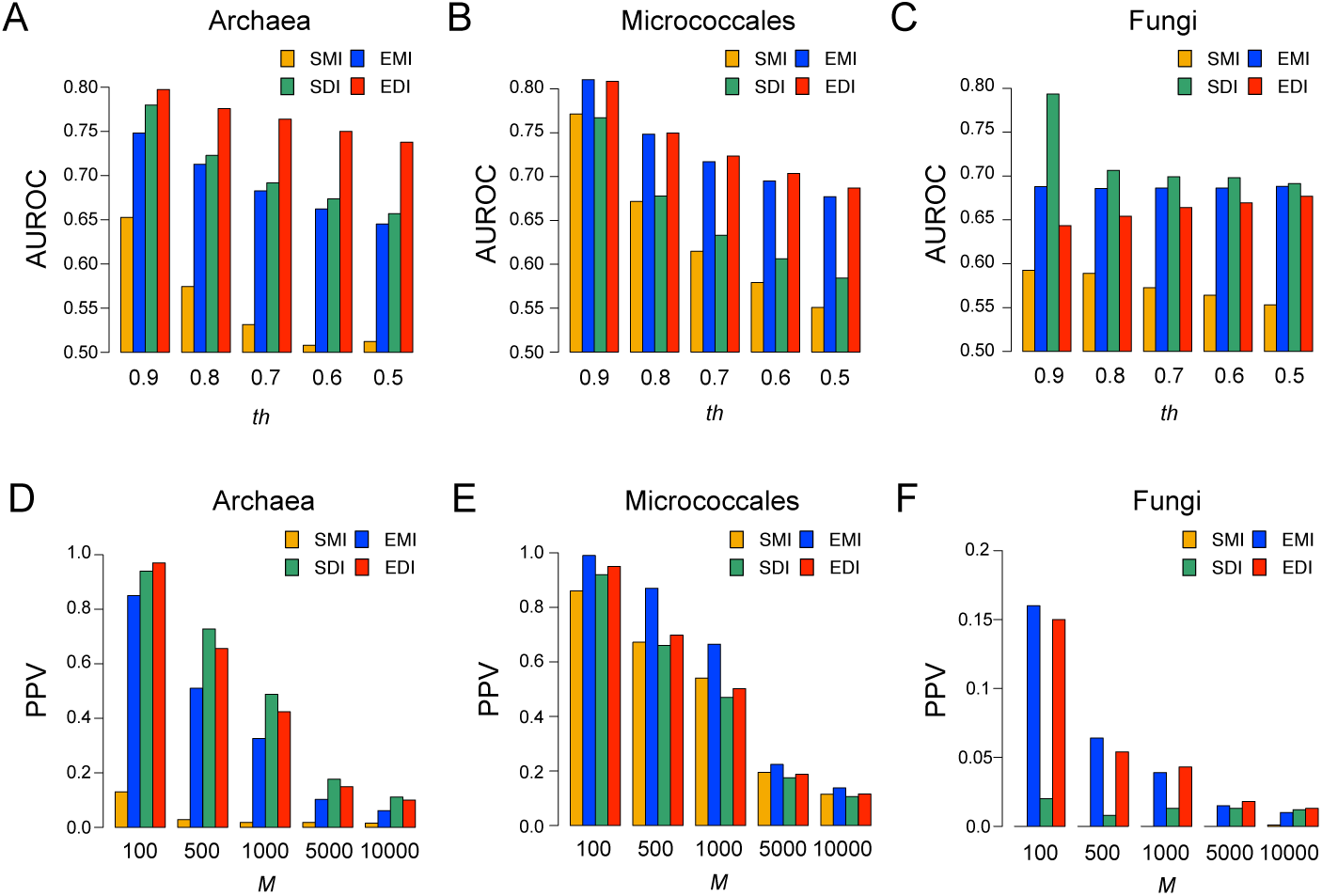
(A-C) Overall discrimination performances of each evaluation measure using the AUC scores. The x-axis represents the *th* value, which defines positive dataset. The y-axis represents the AUROC score. (A), (B), and (C) panels represent results for the archaea, micrococcales, and fungi datasets, respectively. (D-F) Prediction performances for highly ranked OG pairs of each evaluation measure (*th* = 0.7). The x-axis represents the *M* value. The y-axis represents the PPV. (D), (E), and (F) panels represent results for the archaea, micrococcales, and fungi datasets, respectively. The yellow, blue, green, and red colors represent SMI, EMI, SDI, and EDI, respectively.

We second investigated the prediction accuracies of highly ranked (top *M*) OG pairs for each evaluation measure (Fig. 1D-F). In almost all cases, SMI exhibited the worst or near-worst performance. On the other hand, the best-performing evaluation measures depended on the datasets and *M*. For example, when *M* was 1,000, SDI, EMI, and EDI showed the highest PPVs for the archaea, micrococcales, and fungi datasets, respectively. Thus, the reconstruction of gene content history and consideration of spurious correlations generally increases performances, which is better and whether the consideration of both performs better depends on the case, consistent with the AUC score analysis.

### Performances of integrated evaluation measures

Because highly ranked OG pairs estimated by EMI, SDI, and EDI showed the best performance depending on the conditions, we next investigated whether their integration showed better performance. There are four combination types for the integration: EMI and SDI, EMI and EDI, SDI and EDI, and all three evaluation measures. For the integration, we first ordered the OG pairs in descending order by thier scores for EMI, SDI, and EDI. Then, for each combination, we sorted the OG pairs by any of the integration types that are the maximum, average or minimum values of their ranks in all measures under consideration.

We investigated the AUC and PPV performances of 12 integrated evaluation measures comprising four combination types and three integration types (Table S8-S13). We found that the best condition for the integrated evaluation measures depended on the dataset and the threshold (*th* or *M*). As a general trend, the integration by the minimum and average values showed the highest score in the AUC and PPV analyses, respectively. In addition, we found that the highest integrated evaluation measures performed better than the highest single evaluation measures, except in the case of the PPV analysis of the micrococcales dataset when *M* = 100, 500 or 10, 000 (Fig. 2). These results strongly suggest that while EMI, SDI, and EDI are good at considering gene content evolutionary history and/or ignoring spurious correlation, they also lose useful information in functional estimation in its own way, which could be salvaged by their integration.

**Figure 2:**
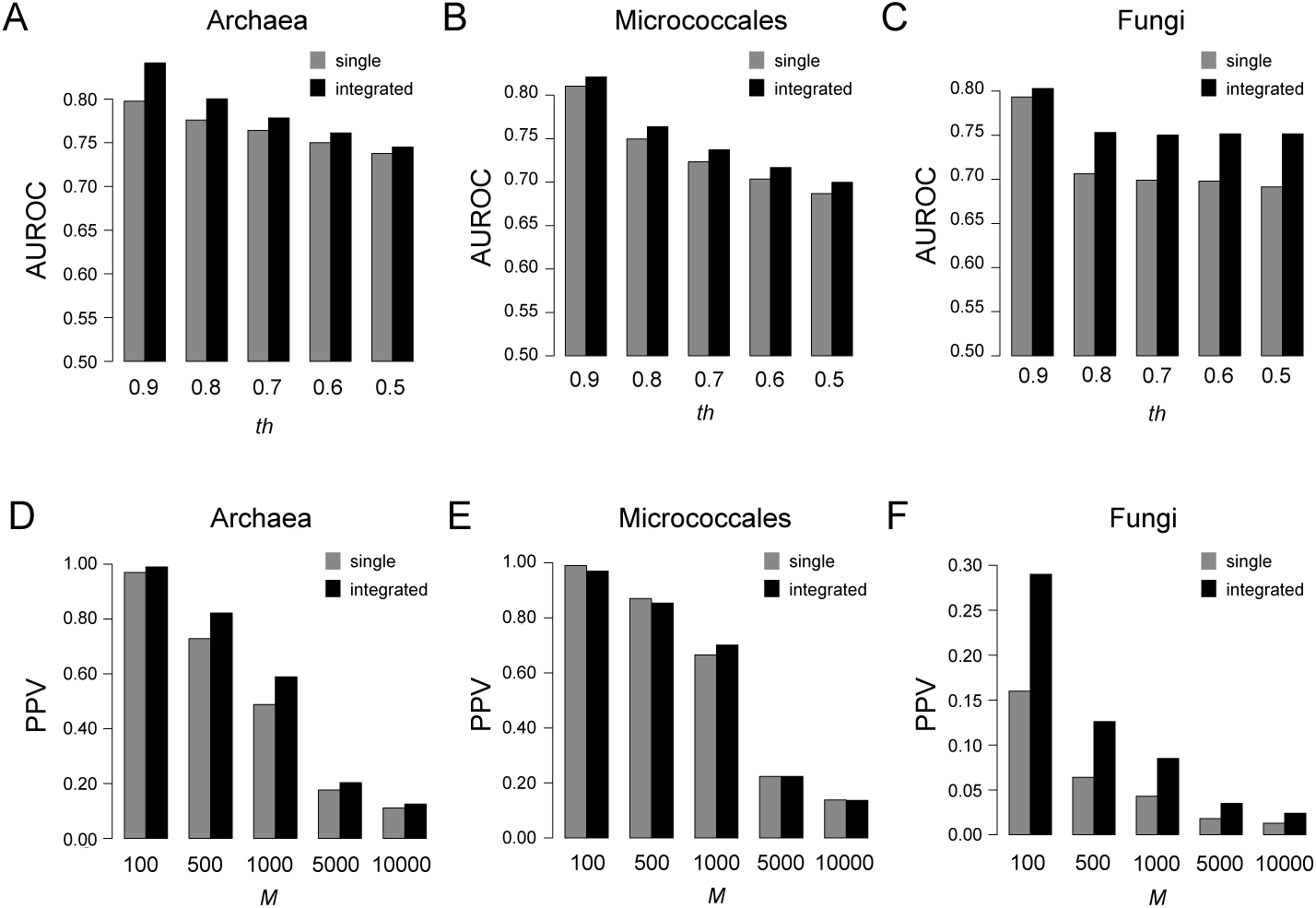
(A-C) Overall discrimination performances of integrated evaluation measures using the AUC scores. The x-axis represents the *th* value, which defines positive dataset. The y-axis represents the AUROC score. (A), (B), and (C) panels represent results for the archaea, micrococcales, and fungi datasets, respectively. (D-F) Prediction performances for highly ranked OG pairs of integrated evaluation measures (*th* = 0.7). The x-axis represents the *M* value. The y-axis represents the PPV. (D), (E), and (F) panels represent results for the archaea, micrococcales, and fungi datasets, respectively. The gray and black colors represent the highest single evaluation measure and integrated evaluation measure, respectively.

### Examples of the detected OG pairs

Finally, as examples of the highly ranked OG pairs, we show lists of the top five ranked OG pairs by the integration of all three evaluation measures (Table 1). We used the average value as the integration type and regarded the value as the prediction score. Except for two cases, these OG pairs had recalculated STRING association scores above 0.9, which means that functional associations had the highest confidence. Most of these gene pairs had known functional relationships. For example, the first rank in the archaea dataset was a pair of *ZnuA* and *ZnuB*, which are components of the ABC-type zinc uptake system. As another example, the fifth rank in the micrococcales dataset was a pair of *DnaC*, which is involved in DNA replication, and COG4584, a transposase.

**Table 1:** The lists of the top 5 OG pairs detected by the combination of all three evaluation measures.

| Taxonomy | Rank | OG1 | OG2 | Prediction score | STRING score |
| --- | --- | --- | --- | --- | --- |
| Archaea | 1 | COG0803 (*ZnuA*) | COG1108 (*ZnuB*) | 10.0 | 0.992 |
|  | 2 | COG1203 (*Cas3*) | COG1688 (*Cas5*) | 14.7 | 0.996 |
|  | 3 | COG1108 (*ZnuB*) | COG1121 (*ZnuC*) | 17.3 | 0.994 |
|  | 4 | COG2998 (*TupA*) | COG4662 (*TupA*) | 21.3 | 0.999 |
|  | 5 | COG1336 (*Cmr4*) | COG1604 (*Cmr6*) | 24.0 | 0.999 |
| Micrococcales | 1 | COG3181 (*TctC*) | COG3333 (*TctA*) | 1.7 | 0.989 |
|  | 2 | COG1135 (*AbcC*) | COG2011 (*MetP*) | 7.0 | 0.995 |
|  | 3 | COG1464 (*NlpA*) | COG2011 COG2011 (*MetP*) | 10.7 | 0.996 |
|  | 4 | COG1135 (*AbcC*) | COG1464 (*NlpA*) | 12.3 | 0.987 |
|  | 5 | COG1484 (*DnaC*) | COG4584 | 12.7 | 0.986 |
| Fungi | 1 | COG0043 (*UbiD*) | COG0163 (*UbiX*) | 34.3 | 0.998 |
|  | 2 | KOG4501 | NOG13474 | 143.3 | 0.0 |
|  | 3 | COG5441 | COG5564 | 620.3 | 0.988 |
|  | 4 | COG0843 (*CyoB*) | COG1290 (*QcrB*) | 682.3 | 0.969 |
|  | 5 | COG2051 (*RPS27A*) | KOG3504 | 774.7 | 0.0 |

The first exceptional pair with the recalculated STRING score of 0.0 was KOG4501 and NOG13473, which was ranked second in the fungi dataset. As explained previously, the recalculated STRING scores depend on gene neighborhood conservation, gene fusion, co-expression, protein interaction experiments, other databases, and text mining; a 0.0 score means that the two OGs have no signals of relationships in any of them. Among them, gene neighborhood conservation and gene fusion are not effective for eukaryotic genome analysis. It is also not surprising that other databases and text mining provide no information, because NOG13473 is a function-unknown gene. Because genes with functional relationships without co-expression and direct interactions are not rare in life systems (such as proteins that function at different time points of the same biological process), we speculated that KOG4501 and NOG13473 may have true functional relationships. The human gene belonging to KOG4501 is *ASCC2*, which is involved in DNA damage repair ([35]). Thus, we argued that NOG13473 may have a DNA damage repair function at an earlier or later stage than that of KOG4501 without direct interaction. In addition, the second exceptional pair was COG2051 and KOG3504, which was ranked fifth in the fungi dataset. Because both these OGs are ribosomal proteins, the recalculated STRING score may suggest the insufficient annotation.

## Discussion

In this study, we evaluated the effectiveness of IPM in the phylogenetic profiling analysis. We constructed four evaluation measures, SMI, EMI, SDI, and EDI, based on whether evolutionary information and the IPM were used. We then investigated the performance of the four evaluation measures using the STRING datasets. We showed that SDI and EDI had the best performances in many cases. In addition, we revealed that predictions based on the combinations of EMI, SDI, and EDI showed higher performance than predictions based on a single evaluation measure. These results demonstrated that the IPM is a powerful approach in phylogenetic profiling.

Although even simple combinations of the evaluation measures yielded good prediction results, more sophisticated methods of combining the evaluation measures may provide better prediction results, for example, machine learning methods. A similar concept was proposed in studies on protein structure prediction based on IPM ([36, 37]). These studies integrated various scores, such as co-evolutionary information using IPM, and predicted solvent accessibility information using supervised machine learning methods, such as deep learning.

We assumed that the input phylogenetic tree and gene content evolutionary history were correct when calculating EMI and EDI. However, they were estimations and intrinsically subject to uncertainty. Such uncertainty should decrease the accuracy of phylogenetic profiling analysis in general ([38]). One solution is to consider the distribution of the estimates by calculating the expected values (instead of counts) of gene gains and losses for each phylogenetic branch. Cohen *et al*. adopted this approach ([11, 39]), but a comparison with other methods has not been conducted and further studies are required. Because this extension requires the use of continuous data, the Gaussian graphical model will need to be used for considering spurious correlations, instead of the Potts model for categorical data ([40]).

In this study, we analyzed only the relationships between two OGs; however, many OGs have higher-order functional relationships among three or more OGs (such as multi-protein complexes). Several studies have focused on the logic relationships of three OGs in phylogenetic profiling ([41, 42, 43]). An example of a logic relationship is C = A ∧ B for OGs A, B, and C, which means that OG C needs both OGs A and B for its function. To date, logic relationship analysis in phylogenetic profiling used local evaluation measures, thus the detection of such higher-order functional relationships based on global evaluation measures is an essential future task. Technically, it is not difficult to extend the Potts model to include (more than) ternary relationships ([44]), but efficient parameter estimation and construction of large-scale datasets for precise parameter estimation will be difficult.

## Acknowledgments

Computations in this research were performed using the supercomputing facilities at the National Institute of Genetics in Research Organization of Information and Systems.

