## Supplementary Materials for "Inverse Potts model improves accuracy of phylogenetic profiling"

### Supplementary Materials for
### Inverse Potts model improves accuracy of phylogenetic profiling

Tsukasa Fukunaga and Wataru Iwasaki

#### Supplementary Figures

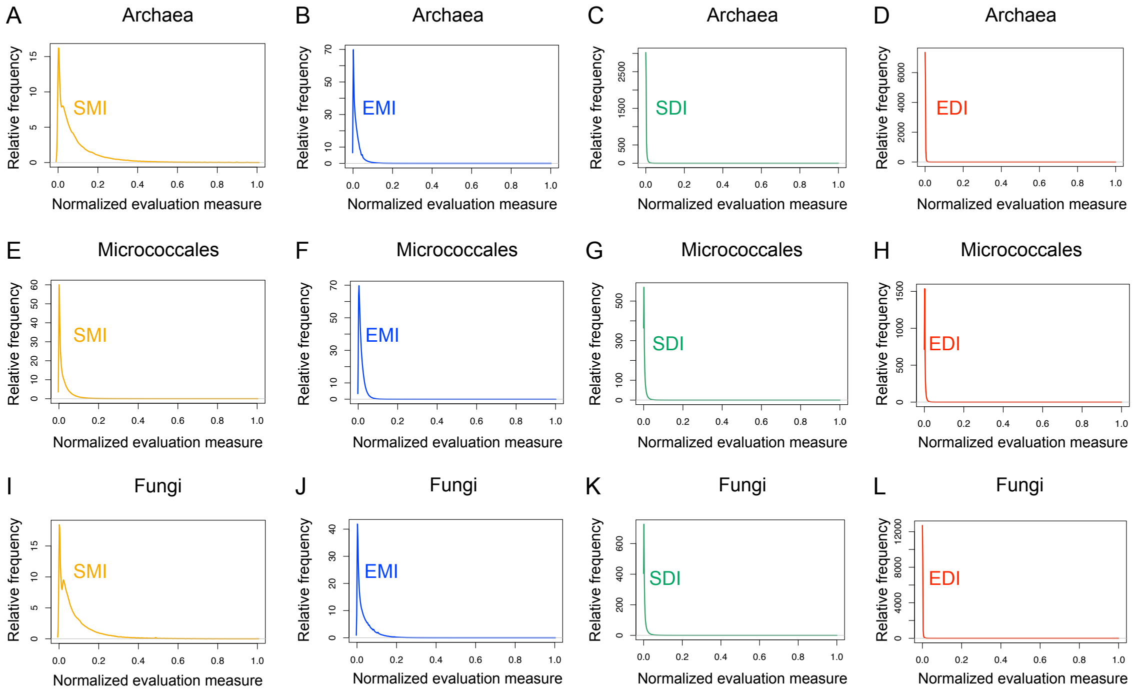


Fig. S1 The distributions of each evaluation measure. The x-axis and the y-axis represent the normalized evaluation measure and the relative frequency, respectively. The yellow, blue, green, and red colors represent the SMI, the EMI, the SDI, and the EDI, respectively. Distributions of (A) the SMI, (B) the EMI, (C) the SDI, and (D) the EDI for the archaea dataset are shown. In addition, distributions of (E) the SMI, (F) the EMI, (G) the SDI, and (H) the EDI for the micrococcales dataset are shown. Furthermore, distributions of (I) the SMI, (J) the EMI, (K) the SDI, and (L) the EDI for the fungi dataset are shown.

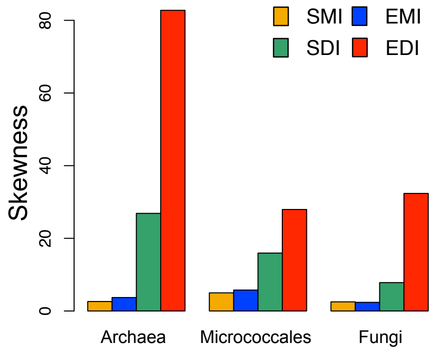


Fig. S2 The skewnesses of distributions of each evaluation measure. The yellow, blue, green, and red colors represent the SMI, the EMI, the SDI and the EDI, respectively. The y-axis represents the skewness.

#### Supplementary Tables

Table S1 The dataset size for each dataset

| taxonomic group | threshold | positive data | negative data |
|---|---|---|---|
| Archaea | 0.9 | 2912 | 4128463 |
| | 0.8 | 7548 | 4123827 |
| | 0.7 | 15112 | 4116263 |
| | 0.6 | 26416 | 4104959 |
| | 0.5 | 46717 | 4084658 |
| Micrococcales | 0.9 | 2426 | 1811134 |
| | 0.8 | 5567 | 1807993 |
| | 0.7 | 9953 | 1803607 |
| | 0.6 | 16666 | 1796894 |
| | 0.5 | 29135 | 1784425 |
| Fungi | 0.9 | 3101 | 16732904 |
| | 0.8 | 6415 | 16729590 |
| | 0.7 | 8968 | 16727037 |
| | 0.6 | 12668 | 16723337 |
| | 0.5 | 18871 | 16717134 |

Table S2 The AUC scores of SDI with varying parameters in the archaea data set

| ($\lambda$, $\epsilon$) | (0.0, 0.01) | (0.0, 0.001) | (0.01, 0.01) | (0.01, 0.001) | (0.05, 0.01) | (0.05, 0.001) | (0.1, 0.01) | (0.1, 0.001) |
|---|---|---|---|---|---|---|---|---|
| $th = 0.9$ | 0.7792 | 0.7785 | 0.7782 | 0.7784 | 0.7774 | 0.7787 | 0.7769 | **0.7799** |
| $th = 0.8$ | **0.7228** | 0.7097 | 0.7225 | 0.7097 | 0.7204 | 0.7100 | 0.7169 | 0.7112 |
| $th = 0.7$ | **0.6918** | 0.6708 | 0.6917 | 0.6708 | 0.6878 | 0.6709 | 0.6819 | 0.6718 |
| $th = 0.6$ | 0.6735 | 0.6511 | **0.6736** | 0.6510 | 0.6687 | 0.6511 | 0.6627 | 0.6518 |
| $th = 0.5$ | **0.6568** | 0.6327 | **0.6568** | 0.6326 | 0.6518 | 0.6324 | 0.6454 | 0.6333 |

| ($\lambda$, $\epsilon$) | (0.5, 0.01) | (0.5, 0.001) | (1.0, 0.01) | (1.0, 0.001) | (5.0, 0.01) | (5.0, 0.001) |
|---|---|---|---|---|---|---|
| $th = 0.9$ | 0.7796 | 0.7790 | 0.7797 | 0.7759 | 0.7489 | 0.7498 |
| $th = 0.8$ | 0.7072 | 0.7061 | 0.6980 | 0.6957 | 0.6610 | 0.6609 |
| $th = 0.7$ | 0.6667 | 0.6656 | 0.6575 | 0.6541 | 0.6255 | 0.6256 |
| $th = 0.6$ | 0.6469 | 0.6459 | 0.6389 | 0.6352 | 0.6078 | 0.6075 |
| $th = 0.5$ | 0.6280 | 0.6265 | 0.6189 | 0.6151 | 0.5889 | 0.5883 |

The bold values are the highest scores in each column.

Table S3 The AUC scores of SDI with varying parameters in the micrococcales data set.

| ($\lambda$, $\epsilon$) | (0.0, 0.01) | (0.0, 0.001) | (0.01, 0.01) | (0.01, 0.001) | (0.05, 0.01) | (0.05, 0.001) | (0.1, 0.01) | (0.1, 0.001) |
|---|---|---|---|---|---|---|---|---|
| $th = 0.9$ | 0.7529 | 0.7614 | 0.7550 | 0.7614 | 0.7613 | 0.7624 | 0.7630 | 0.7631 |
| $th = 0.8$ | 0.6723 | 0.6734 | 0.6746 | 0.6734 | **0.6779** | 0.6739 | 0.6767 | 0.6740 |
| $th = 0.7$ | 0.6278 | 0.6267 | 0.6306 | 0.6266 | **0.6330** | 0.6267 | 0.6308 | 0.6265 |
| $th = 0.6$ | 0.6020 | 0.5969 | 0.6047 | 0.5969 | **0.6063** | 0.5966 | 0.6030 | 0.5962 |
| $th = 0.5$ | 0.5809 | 0.5742 | 0.5837 | 0.5742 | **0.5844** | 0.5740 | 0.5807 | 0.5734 |

| ($\lambda$, $\epsilon$) | (0.5, 0.01) | (0.5, 0.001) | (1.0, 0.01) | (1.0, 0.001) | (5.0, 0.01) | (5.0, 0.001) |
|---|---|---|---|---|---|---|
| $th = 0.9$ | **0.7669** | 0.7668 | 0.7616 | 0.7619 | 0.7255 | 0.7266 |
| $th = 0.8$ | 0.6734 | 0.6733 | 0.6643 | 0.6646 | 0.6309 | 0.6297 |
| $th = 0.7$ | 0.6229 | 0.6226 | 0.6126 | 0.6123 | 0.5868 | 0.5861 |
| $th = 0.6$ | 0.5900 | 0.5892 | 0.5788 | 0.5789 | 0.5607 | 0.5603 |
| $th = 0.5$ | 0.5657 | 0.5650 | 0.5551 | 0.5553 | 0.5434 | 0.543 |

The bold values are the highest scores in each column.

Table S4 The AUC scores of SDI with varying parameters in the fungi data set.

| ($\lambda$, $\epsilon$) | (0.0, 0.01) | (0.0, 0.001) | (0.01, 0.01) | (0.01, 0.001) | (0.05, 0.01) | (0.05, 0.001) | (0.1, 0.01) | (0.1, 0.001) | (0.5, 0.01) | (0.5, 0.001) | (1.0, 0.01) | (1.0, 0.001) | (5.0, 0.01) | (5.0, 0.001) |
|---|---|---|---|---|---|---|---|---|---|---|---|---|---|---|
| $th = 0.9$ | 0.7308 | 0.7295 | 0.7523 | 0.7391 | 0.7434 | 0.7292 | 0.7397 | 0.7346 | 0.7697 | 0.7359 | **0.7934** | 0.7477 | 0.7245 | 0.7709 |
| $th = 0.8$ | 0.6565 | 0.6498 | 0.6721 | 0.6579 | 0.6697 | 0.6515 | 0.6671 | 0.6563 | 0.6882 | 0.6586 | **0.7064** | 0.6643 | 0.6515 | 0.6701 |
| $th = 0.7$ | 0.6518 | 0.6428 | 0.6667 | 0.6506 | 0.6651 | 0.6443 | 0.6630 | 0.6489 | 0.6783 | 0.6474 | **0.6991** | 0.6524 | 0.6462 | 0.6553 |
| $th = 0.6$ | 0.6499 | 0.6406 | 0.6660 | 0.6477 | 0.6629 | 0.6416 | 0.6609 | 0.6457 | 0.6755 | 0.6427 | **0.6981** | 0.6481 | 0.6480 | 0.6535 |
| $th = 0.5$ | 0.6483 | 0.6367 | 0.6631 | 0.6433 | 0.6601 | 0.6375 | 0.6582 | 0.6408 | 0.6687 | 0.6338 | **0.6914** | 0.6366 | 0.6472 | 0.6416 |

The bold values are the highest scores in each column.

Table S5 The AUC scores of EDI with varying parameters in the archaea data set.

| ($\lambda$, $\epsilon$) | (0.0, 0.01) | (0.0, 0.001) | (0.01, 0.01) | (0.01, 0.001) | (0.05, 0.01) | (0.05, 0.001) | (0.1, 0.01) | (0.1, 0.001) | (0.5, 0.01) | (0.5, 0.001) | (1.0, 0.01) | (1.0, 0.001) | (5.0, 0.01) | (5.0, 0.001) |
|---|---|---|---|---|---|---|---|---|---|---|---|---|---|---|
| $th = 0.9$ | 0.7954 | 0.7573 | **0.7974** | 0.7566 | 0.7901 | 0.7570 | 0.7804 | 0.7560 | 0.7374 | 0.7341 | 0.6998 | 0.6993 | 0.5935 | 0.6005 |
| $th = 0.8$ | 0.7751 | 0.7307 | **0.7759** | 0.7305 | 0.7669 | 0.7310 | 0.7555 | 0.7305 | 0.7140 | 0.7094 | 0.6732 | 0.6750 | 0.5606 | 0.5655 |
| $th = 0.7$ | **0.7640** | 0.7118 | 0.7635 | 0.7119 | 0.7523 | 0.7122 | 0.7391 | 0.7117 | 0.6950 | 0.6915 | 0.6542 | 0.6555 | 0.5367 | 0.5398 |
| $th = 0.6$ | **0.7502** | 0.6950 | 0.7493 | 0.6952 | 0.7374 | 0.6957 | 0.7234 | 0.6954 | 0.6781 | 0.6751 | 0.6363 | 0.6379 | 0.5171 | 0.5207 |
| $th = 0.5$ | **0.7378** | 0.6819 | 0.7365 | 0.6819 | 0.7249 | 0.6824 | 0.7114 | 0.6822 | 0.6640 | 0.6611 | 0.6210 | 0.6228 | 0.5061 | 0.5083 |

The bold values are the highest scores in each column.

Table S6 The AUC scores of EDI with varying parameters in the micrococcales data set.

| ($\lambda$, $\epsilon$) | (0.0, 0.01) | (0.0, 0.001) | (0.01, 0.01) | (0.01, 0.001) | (0.05, 0.01) | (0.05, 0.001) | (0.1, 0.01) | (0.1, 0.001) |
|---|---|---|---|---|---|---|---|---|
| $th = 0.9$ | 0.7941 | 0.8043 | 0.7969 | 0.8044 | 0.8037 | 0.8042 | 0.8085 | **0.8036** |
| $th = 0.8$ | 0.7397 | 0.7434 | 0.7423 | 0.7438 | 0.7481 | 0.7438 | 0.7497 | **0.7435** |
| $th = 0.7$ | 0.7160 | 0.7172 | 0.7187 | 0.7175 | 0.7233 | **0.7179** | 0.7229 | 0.7180 |
| $th = 0.6$ | 0.6978 | 0.6981 | 0.7005 | 0.6983 | 0.7036 | **0.6988** | 0.7031 | 0.6990 |
| $th = 0.5$ | 0.6818 | 0.6815 | 0.6847 | 0.6817 | 0.6869 | **0.6825** | 0.6859 | 0.6828 |

| ($\lambda$, $\epsilon$) | (0.5, 0.01) | (0.5, 0.001) | (1.0, 0.01) | (1.0, 0.001) | (5.0, 0.01) | (5.0, 0.001) |
|---|---|---|---|---|---|---|
| $th = 0.9$ | 0.7966 | 0.7964 | 0.7811 | 0.7806 | 0.7032 | 0.7119 |
| $th = 0.8$ | 0.7358 | 0.7363 | 0.7201 | 0.7211 | 0.6393 | 0.6391 |
| $th = 0.7$ | 0.7106 | 0.7116 | 0.6985 | 0.6995 | 0.6140 | 0.6167 |
| $th = 0.6$ | 0.6935 | 0.6943 | 0.6828 | 0.6840 | 0.5993 | 0.6016 |
| $th = 0.5$ | 0.6789 | 0.6793 | 0.6699 | 0.6711 | 0.5861 | 0.5886 |

The bold values are the highest scores in each column.

Table S7 The AUC scores of EDI with varying parameters in the fungi data set.

| ($\lambda$, $\epsilon$) | (0.0, 0.01) | (0.0, 0.001) | (0.01, 0.01) | (0.01, 0.001) | (0.05, 0.01) | (0.05, 0.001) | (0.1, 0.01) | (0.1, 0.001) |
|---|---|---|---|---|---|---|---|---|
| $th = 0.9$ | 0.6354 | 0.6194 | **0.6435** | 0.6116 | 0.6432 | 0.6233 | 0.6380 | 0.6253 |
| $th = 0.8$ | 0.6459 | 0.6370 | **0.6541** | 0.6316 | **0.6541** | 0.6404 | 0.6512 | 0.6421 |
| $th = 0.7$ | 0.6568 | 0.6445 | **0.6640** | 0.6394 | 0.6626 | 0.6479 | 0.6589 | 0.6495 |
| $th = 0.6$ | 0.6628 | 0.6451 | **0.6695** | 0.6399 | 0.6671 | 0.6483 | 0.6617 | 0.6499 |
| $th = 0.5$ | 0.6696 | 0.6509 | **0.6769** | 0.6462 | 0.6735 | 0.6540 | 0.6674 | 0.6554 |

| ($\lambda$, $\epsilon$) | (0.5, 0.01) | (0.5, 0.001) | (1.0, 0.01) | (1.0, 0.001) | (5.0, 0.01) | (5.0, 0.001) |
|---|---|---|---|---|---|---|
| $th = 0.9$ | 0.5946 | 0.5911 | 0.5513 | 0.5510 | 0.5341 | 0.5425 |
| $th = 0.8$ | 0.6222 | 0.6203 | 0.5898 | 0.5916 | 0.4934 | 0.5081 |
| $th = 0.7$ | 0.6310 | 0.6294 | 0.6008 | 0.6028 | 0.5169 | 0.5179 |
| $th = 0.6$ | 0.6311 | 0.6291 | 0.6013 | 0.6031 | 0.5181 | 0.5181 |
| $th = 0.5$ | 0.6379 | 0.6361 | 0.6097 | 0.6116 | 0.5246 | 0.5235 |

The bold values are the highest scores in each column.

Table S8 The AUC scores of integrated evaluation measures in the archaea data set.

| | EMI&SDI max | EMI&SDI avg | EMI&SDI min | EMI&EDI max | EMI&EDI avg | EMI&EDI min | SDI&EDI max | SDI&EDI avg |
|---|---|---|---|---|---|---|---|---|
| $th = 0.9$ | 0.7703 | 0.7993 | 0.8027 | 0.7804 | 0.7913 | 0.7858 | 0.7728 | 0.8162 |
| $th = 0.8$ | 0.7297 | 0.7566 | 0.7540 | 0.7535 | 0.7662 | 0.7610 | 0.7405 | 0.7818 |
| $th = 0.7$ | 0.7030 | 0.7264 | 0.7182 | 0.7351 | 0.7463 | 0.7382 | 0.7220 | 0.7630 |
| $th = 0.6$ | 0.6864 | 0.7064 | 0.6938 | 0.7189 | 0.7289 | 0.7192 | 0.7073 | 0.7477 |
| $th = 0.5$ | 0.6709 | 0.6833 | 0.6734 | 0.7048 | 0.7136 | 0.7026 | 0.6935 | 0.7334 |

| | SDI&EDI min | all max | all avg | all min |
|---|---|---|---|---|
| $th = 0.9$ | **0.8412** | 0.7721 | 0.8149 | 0.8267 |
| $th = 0.8$ | **0.8002** | 0.7382 | 0.7839 | 0.7873 |
| $th = 0.7$ | **0.7783** | 0.7177 | 0.7623 | 0.7580 |
| $th = 0.6$ | **0.7611** | 0.7026 | 0.7448 | 0.7364 |
| $th = 0.5$ | **0.7451** | 0.6883 | 0.7288 | 0.7172 |

The bold values are the highest scores in each column.

Table S9 The AUC scores of integrated evaluation measures in the micrococcales data set.

| | EMI&SDI max | EMI&SDI avg | EMI&SDI min | EMI&EDI max | EMI&EDI avg | EMI&EDI min | SDI&EDI max | SDI&EDI avg |
|---|---|---|---|---|---|---|---|---|
| $th = 0.9$ | 0.7918 | 0.8070 | 0.8098 | 0.8040 | 0.8144 | **0.8211** | 0.7850 | 0.8026 |
| $th = 0.8$ | 0.7180 | 0.7396 | 0.7438 | 0.7453 | 0.7574 | **0.7637** | 0.7146 | 0.7371 |
| $th = 0.7$ | 0.6796 | 0.7027 | 0.7066 | 0.7161 | 0.7302 | **0.7372** | 0.6789 | 0.7039 |
| $th = 0.6$ | 0.6554 | 0.6786 | 0.6817 | 0.6949 | 0.7096 | **0.7167** | 0.6551 | 0.6810 |
| $th = 0.5$ | 0.6343 | 0.6583 | 0.6613 | 0.6768 | 0.6923 | **0.6999** | 0.6351 | 0.6614 |

| | SDI&EDI min | all max | all avg | all min |
|---|---|---|---|---|
| $th = 0.9$ | 0.8065 | 0.7906 | 0.8161 | 0.8208 |
| $th = 0.8$ | 0.7433 | 0.7211 | 0.7553 | 0.7591 |
| $th = 0.7$ | 0.7106 | 0.6846 | 0.7241 | 0.7260 |
| $th = 0.6$ | 0.6880 | 0.6608 | 0.7019 | 0.7028 |
| $th = 0.5$ | 0.6686 | 0.6400 | 0.6830 | 0.6836 |

The bold values are the highest scores in each column.

Table S10 The AUC scores of integrated evaluation measures in the fungi data set.

| | EMI&SDI max | EMI&SDI avg | EMI&SDI min | EMI&EDI max | EMI&EDI avg | EMI&EDI min | SDI&EDI max | SDI&EDI avg |
|---|---|---|---|---|---|---|---|---|
| $th = 0.9$ | 0.7296 | 0.7822 | 0.7930 | 0.6514 | 0.6791 | 0.6922 | 0.7024 | 0.7636 |
| $th = 0.8$ | 0.6927 | 0.7340 | 0.7445 | 0.6570 | 0.6833 | 0.6957 | 0.6669 | 0.7204 |
| $th = 0.7$ | 0.6918 | 0.7312 | 0.7391 | 0.6649 | 0.6890 | 0.6992 | 0.6703 | 0.7221 |
| $th = 0.6$ | 0.6924 | 0.7315 | 0.7381 | 0.6674 | 0.6921 | 0.7028 | 0.6727 | 0.7247 |
| $th = 0.5$ | 0.6917 | 0.7300 | 0.7357 | 0.6734 | 0.6973 | 0.7072 | 0.6743 | 0.7252 |

| | SDI&EDI min | all max | all avg | all min |
|---|---|---|---|---|
| $th = 0.9$ | **0.8029** | 0.6954 | 0.7575 | 0.7917 |
| $th = 0.8$ | 0.7504 | 0.6641 | 0.7279 | **0.7531** |
| $th = 0.7$ | 0.7488 | 0.6678 | 0.7299 | **0.7502** |
| $th = 0.6$ | 0.7512 | 0.6695 | 0.7323 | **0.7515** |
| $th = 0.5$ | 0.7506 | 0.6718 | 0.7344 | **0.7515** |

The bold values are the highest scores in each column.

Table S11 The PPV scores of integrated evaluation measures in the archaea data set.

| | EMI&SDI max | EMI&SDI avg | EMI&SDI min | EMI&EDI max | EMI&EDI avg | EMI&EDI min | SDI&EDI max | SDI&EDI avg |
|---|---|---|---|---|---|---|---|---|
| $M = 100$ | 0.980 | 0.980 | 0.960 | **0.990** | **0.990** | 0.960 | 0.980 | 0.970 |
| $M = 500$ | 0.788 | 0.790 | 0.714 | 0.766 | 0.776 | 0.656 | 0.768 | 0.778 |
| $M = 1000$ | 0.576 | 0.583 | 0.489 | 0.517 | 0.530 | 0.445 | 0.514 | 0.527 |
| $M = 5000$ | 0.180 | 0.186 | 0.164 | 0.174 | 0.180 | 0.147 | 0.180 | 0.183 |
| $M = 10000$ | 0.108 | 0.114 | 0.102 | 0.109 | 0.112 | 0.092 | 0.112 | 0.115 |

| | SDI&EDI min | all max | all avg | all min |
|---|---|---|---|---|
| $M = 100$ | 0.970 | **0.990** | **0.990** | 0.970 |
| $M = 500$ | 0.748 | 0.804 | **0.822** | 0.738 |
| $M = 1000$ | 0.491 | 0.569 | **0.589** | 0.503 |
| $M = 5000$ | 0.176 | 0.193 | **0.203** | 0.173 |
| $M = 10000$ | 0.118 | 0.116 | **0.125** | 0.109 |

The bold values are the highest scores in each column.

Table S12 The PPV scores of integrated evaluation measures in the micrococcales data set.

| | EMI&SDI max | EMI&SDI avg | EMI&SDI min | EMI&EDI max | EMI&EDI avg | EMI&EDI min | SDI&EDI max | SDI&EDI avg |
|---|---|---|---|---|---|---|---|---|
| $M = 100$ | 0.950 | 0.950 | 0.950 | 0.950 | 0.960 | **0.970** | 0.950 | 0.950 |
| $M = 500$ | 0.850 | 0.844 | **0.854** | 0.790 | 0.806 | 0.820 | 0.808 | 0.800 |
| $M = 1000$ | 0.687 | **0.701** | 0.682 | 0.637 | 0.645 | 0.656 | 0.661 | 0.668 |
| $M = 5000$ | 0.213 | 0.221 | **0.224** | 0.204 | 0.208 | 0.222 | 0.196 | 0.200 |
| $M = 10000$ | 0.125 | 0.131 | 0.136 | 0.121 | 0.124 | **0.137** | 0.115 | 0.117 |

| | SDI&EDI min | all max | all avg | all min |
|---|---|---|---|---|
| $M = 100$ | 0.950 | 0.950 | 0.950 | 0.950 |
| $M = 500$ | 0.732 | 0.840 | 0.848 | 0.828 |
| $M = 1000$ | 0.516 | 0.685 | 0.696 | 0.643 |
| $M = 5000$ | 0.194 | 0.210 | 0.215 | 0.217 |
| $M = 10000$ | 0.117 | 0.124 | 0.129 | **0.137** |

The bold values are the highest scores in each column.

Table S13 The PPV scores of integrated evaluation measures in the fungi data set.

| | EMI&SDI max | EMI&SDI avg | EMI&SDI min | EMI&EDI max | EMI&EDI avg | EMI&EDI min | SDI&EDI max | SDI&EDI avg |
|---|---|---|---|---|---|---|---|---|
| $M = 100$ | 0.120 | 0.250 | 0.190 | 0.190 | 0.220 | 0.230 | 0.120 | 0.160 |
| $M = 500$ | 0.056 | **0.126** | 0.120 | 0.070 | 0.088 | 0.092 | 0.042 | 0.090 |
| $M = 1000$ | 0.035 | **0.085** | 0.076 | 0.042 | 0.067 | 0.063 | 0.029 | 0.060 |
| $M = 5000$ | 0.016 | **0.035** | 0.034 | 0.018 | 0.023 | 0.023 | 0.019 | 0.026 |
| $M = 10000$ | 0.013 | **0.024** | 0.023 | 0.014 | 0.016 | 0.016 | 0.016 | 0.016 |

| | SDI&EDI min | all max | all avg | all min |
|---|---|---|---|---|
| $M = 100$ | 0.130 | 0.150 | **0.290** | **0.290** |
| $M = 500$ | 0.082 | 0.060 | 0.120 | 0.106 |
| $M = 1000$ | 0.062 | 0.038 | 0.081 | 0.074 |
| $M = 5000$ | 0.024 | 0.019 | 0.031 | 0.029 |
| $M = 10000$ | 0.018 | 0.015 | 0.021 | 0.019 |

The bold values are the highest scores in each column.
